# Population divergence time estimation using individual lineage label switching

**DOI:** 10.1101/587832

**Authors:** Peter Beerli, Haleh Ashki, Somayeh Mashayekhi, Michal Palczewski

## Abstract

Divergence time estimation from multilocus genetic data has become common in population genetics and phylogenetics. We present a new Bayes inference method that treats the divergence time as a random variable. The divergence time is calculated from an assembly of splitting events on individual lineages in a genealogy. The waiting time for such a splitting event is drawn from a hazard function of the truncated normal distribution. This allows easy integration into the standard coalescence framework used in programs such as MIGRATE. We explore the accuracy of the new inference method with simulated population splittings over a wide range of divergence time values and with a dataset of the Zika virus; the geographic analyses of the expansion of the pathogen follows a trajectory from Africa to Asia to America, corroborating analyses based only on the dates of incidences. Evaluations of simple divergence models show high accuracy, whereas the accuracy of the results of isolation with migration (IM) models depend on the magnitude of the immigration rate and potentially on the number of samples. High immigration rates lead to a time of the most recent common ancestor of the sample that predates the divergence time, thus loses any potential signal of the divergence event in the sample data. This reduced accuracy with high immigration rates is problematic for all IM methods, including ours.

---

In phylogenetics and population genetics often we need to know the time when populations split and evolved independently or when populations started to have reduced gene flow among them; Wakeley and Hey (1997) define an *isolation* model (I) in which the divergence marks the time when the ancestral population split into two groups of individuals that stop exchanging genetic material with each other. Given that most populations within a species may still exchange migrants after divergence, this definition seems overly strict. Consequently, Nielsen and Wakeley (2001) developed the *isolation with migration* model (IM); in their model, the divergence time marks the change from a panmictic ancestral population to two populations linked by gene flow. In both models the divergence times of the populations is always predated by the divergence time of the genes (cf Edwards and Beerli 2000; Arbogast et al. 2002). Both the isolation model and the isolation-with-migration model became popular and were implemented in several software packages, such as BPP (Yang and Rannala 2010), IMa(Hey 2010), Lamarc (Kuhner 2006), and BEAST 2 (Bouckaert et al. 2014).

The isolation with migration model treats the divergence time as a boundary between two models: a structured coalescent population with migration and a panmictic, ancestral population. We describe here an approach that combines migration and divergence within the same structured coalescence framework allowing the boundary to be more fluid. The extent of the boundary is defined by two parameters, the mean of the distribution of the boundary, the divergence time, and the standard deviation of the boundary. We implemented the new method in the program Migrate (Beerli 2006) which was used for all evaluations in this research. The MIT-licensed, open source software Migrate is available from the website http://popgen.sc.fsu.edu.

## Methods

All current coalescence-based methods for estimating a divergence time *τ* between two populations treat the time as a boundary between two different models: the panmictic, ancestral population modeled using the single population coalescent and a population with two subpopulations using the structured coalescent with migration. In a Bayesian inference method, the boundary is adjusted using a prior distribution.

Here we propose a different model. We consider the divergence time as a random variable with a normal distribution. The mean and standard deviation of this distribution are unknown and estimated. We use this distribution to draw times for divergence events for each lineage. We assume that we know the population or species label of the sampled individuals. Looking backward in time, each sample lineage will be at risk to switch labels irreversibly from a ‘derived’ to an ‘ancestral’ state. Therefore, at a given time in the genealogy, some lineages are in the ancestral population, and some are not. Figure 1 shows an example with a divergence times *τ*. If we assume that this time is fixed, then the figure represents the isolation-with-migration model. If we assume that we have a normal-distributed divergence time with parameters *τ* and *σ_τ_*, then individual lineages can change their state from the ‘descendant’ state to the ‘ancestor’ state by drawing times from this distribution and inserting a ‘divergence’ event. This process is similar to how migration events are drawn.

**Figure 1:**
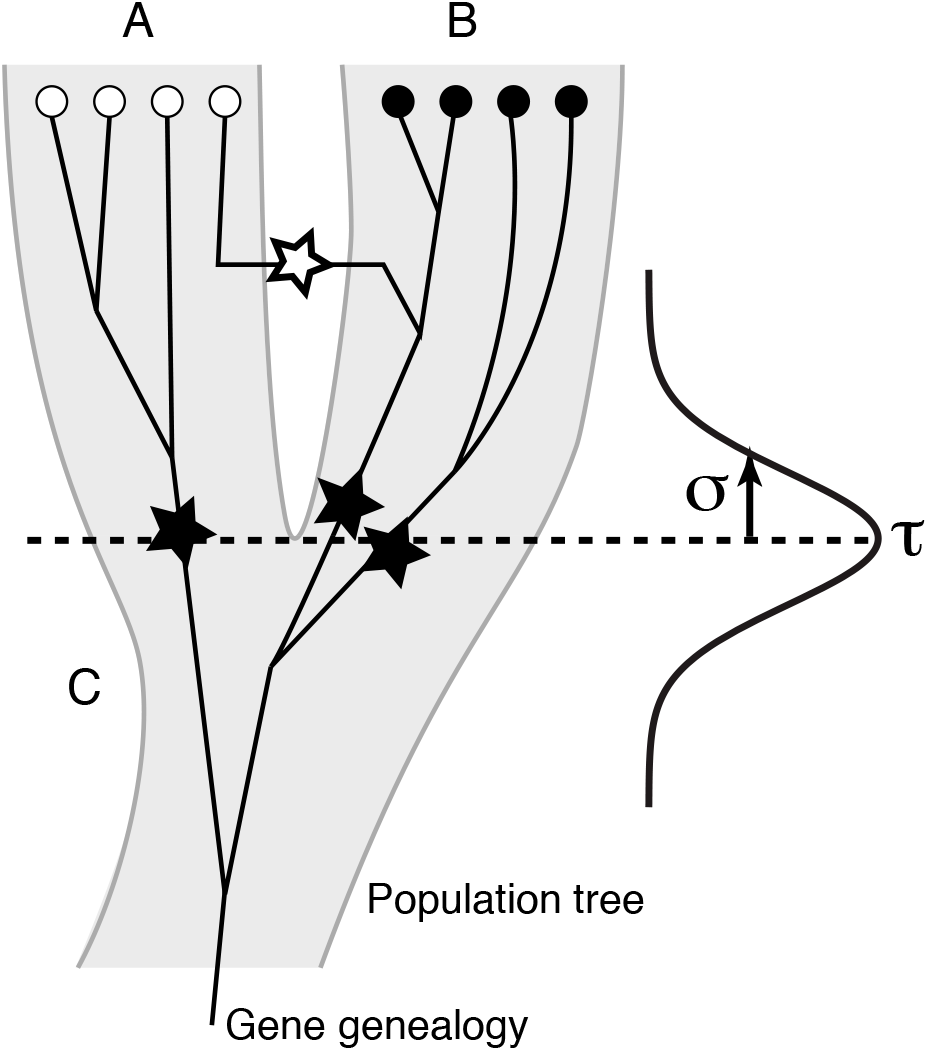
Visualization of population splitting of 2 populations A and B split at times *τ*, lineages in A and B split from the ancestral population C: the divergence time is based on individual lineage population label switching events (dark star) drawn from a distribution with mean (*τ*) and standard deviation (*σ_τ_*); migration events (white star) are drawn from the standard structured coalescent.

### Hazard functions and waiting times

Population genetic inferences commonly use a sample of individuals collected recently, and we are interested in the potential interactions of these individuals in the past. The coalescent (Kingman 1982), allowing probabilistic reconstruction of potential, past genealogies of the sample, leads to inferences of population sizes, migration rates, and other population genetic quantities. For example, in the migration model introduced by Strobeck (1987); Hudson (1991) and used for population parameter inference by Beerli and Felsenstein (2001); Beerli (2006) the times of coalescence events and migration events are drawn from an exponential distribution with a rate that is defined by parameters for population size, migration rate, and the number of sampled lineages in each population. Looking backward in time, a sample lineage is at risk of a coalescence event or a migration event. The risk of coalescence or immigration, switching population labels, in this framework is constant for a given sample size. We can think of this process as a hazard that the lineages will fail to maintain their current status. The *hazard function* is commonly used in an engineering or survival context, were the condition of an object can suddenly change at any given time; the hazard can be constant, increasing, or decreasing with time. The function is expressed as a ratio of the probability density function *f*(·) and the complement of its cumulative density function *F*(·). For the exponential probability density function this hazard is a constant, and for the Kingman coalescent this is

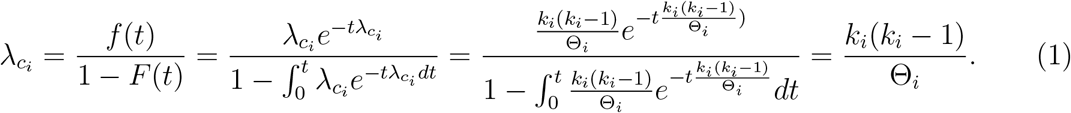

where Θ*_i_* is the mutation-scaled effective population size and *k_i_* is the number of lineages in population *i* at time *t* (Wright 1951; Kingman 1982).

In the structured coalescence, migration is treated similarly. An immigration event happens at the rate *M_ji_* for each lineage, where *M_ji_* is the mutation-scaled immigration rate from population *j* into *i* which is equivalent to the immigration rate *m_ji_* divided by the mutation rate *μ*. The total rate of migration is the sum of all possible migration rates *M_ji_* over all the lineages that have not yet coalesced:

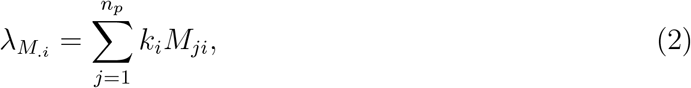

where *n_p_* is the number of populations, and *k_i_* is the number of lineages in population *i* at time *t*. Since the rate of immigrations and coalescences 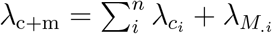 is independent of the waiting time *t*, which shows the time before a coalescent or migration event happens in [*t*_0_, *t*_0_ + *u*], has a probability density function of the exponential mixture

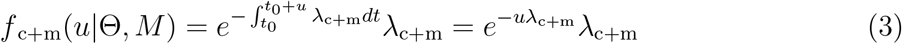

### Divergence time as events on lineages

In 2000 Nielsen and Slatkin introduced and later (Hey and Nielsen 2007) refined a model that adds population splitting, thus removing the assumption that populations are present for a very long time without removing the assumption of migration between the populations. We have developed an alternative to Nielsen’s and Hey’s approach that allows distributing the analysis onto cluster computers and using large datasets.

We treat the time of splitting as a random variable with a particular probability density. We chose to use the zero-truncated normal distribution because it has two parameters: mean and standard deviation. These parameters are commonly used and discuss quantities of interest. The mean describes the most likely time of the population divergence and the standard deviation describes the uncertainty of that divergence time. We consider the truncated normal distribution a good choice to discuss divergence times, but it certainly is not the only possible distribution. We could have used a Weibull distribution or a Gamma distribution, both have a natural bound at zero, but their standard parameters are less familiar to biologists.

Looking backward in time, we know the fate of a lineage sampled today; any individual sampled today must have started in an ancestral population; thus, each lineage sampled today is at risk to switch from the derived to the ancestral population. We assume that the risk of failure to stay in the derived population is increasing the further back in time the process moves. This process can be expressed with a hazard function of the normal distribution that is not constant, in contrast to the hazard function of the exponential distribution. The use of this hazard function allows us to integrate our population splitting distribution into our coalescence with migration framework. To express the risk of switching the population label (population splitting) we calculate the rate of splitting (divergence) events 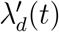 by using the hazard function of a truncated normal distribution with bounds *b*_0_ = 0.0 and *b*_1_ as

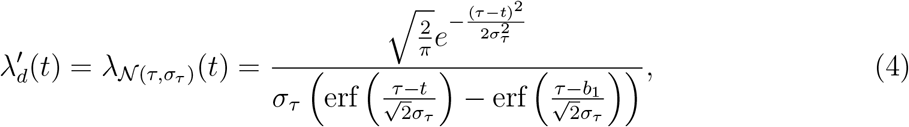

*τ* and *σ_τ_* are the parameters of the normal distribution; erf is the error function

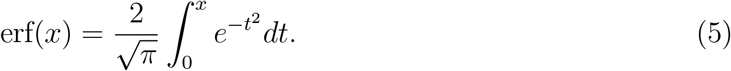

To calculate the probability that no splitting event happens in the interval [*t*_0_, *t*_0_ + *u*] we integrate and get

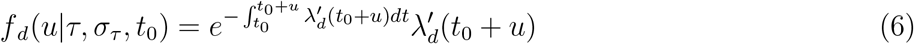

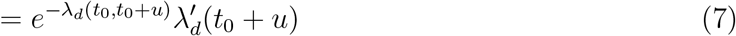

where

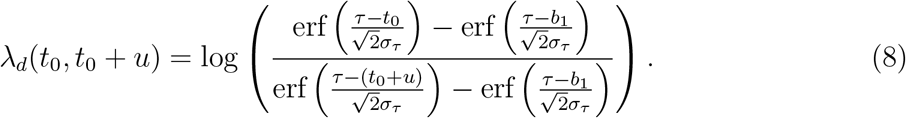

Combining these individual waiting times for coalescence, immigration, and splitting leads to the overall probability density for the waiting time *u* to the next event in the interval [*t*_0_, *t*_0_ + *u*]

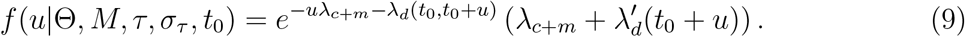

### Genealogy-probability calculations

The posterior density distribution *p*(*ρ*|*D*) for all parameters *ρ*, such as mutation-scaled population size Θ, mutation-scaled immigration rates *M*, divergence time mean *τ* and standard deviation *σ_τ_* is

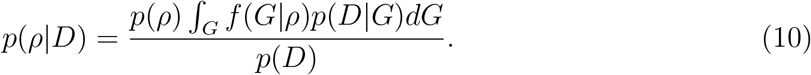

We approximate the posterior distributions with histograms for each parameter *ρ* collected through the Markov chain Monte Carlo run. The Metropolis-Hastings acceptance/rejection steps will need calculations of the likelihood of the genealogy *p*(*D*|*G*) and the parameters *p*(*G*|*ρ*) (Beerli and Felsenstein 1999; Beerli 2006). The likelihood of the genealogy is calculated using the familiar likelihood pruning algorithm used in phylogenetics (Felsenstein 1981). We discuss the evaluation of *p*(*G*|*ρ*) in the next section.

### Probability of events and calculation of the probability density of a genealogy given all parameters

The coalescence process reduces the number of lineages when looking backward in time; coalescent, migration, and divergence events are independent from the events before them. Thus, we can calculate the probability density of a genealogy given all parameters *f*(*G*|*ρ*) as the product over all time intervals

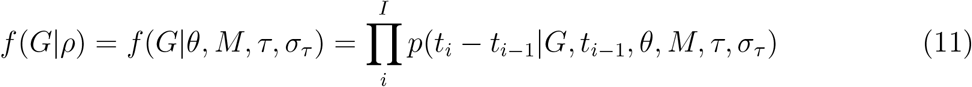

The calculation of *p*(*t_i_* − *t*_*i*−1_|*G*, *t*_*i*−1_, *θ*, *M*, *τ*, *σ_τ_*) is more involved. For each time interval we calculate the exponential waiting time for any event, calculate the probability that the particular event type recorded on the genealogy is drawn, and also need to adjust for how many possible events of the same type can be drawn. If we have a time interval that ends with a coalescent event, then, in the most general case, we calculate

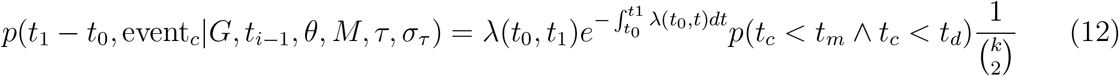

where λ(*t*_0_, *t*_1_) is the sum of all rates for all parameters, for example this includes Eq. (1, 2, 4). There are similar formulae for cases when the interval ends with a divergence event or ends with an immigration event. In a model with only coalescence and migration events this simplifies greatly because *p*(*t_c_* < *t_m_* ∧ *t_c_* < *t_d_*) reduces to *p*(*t_c_* < *t_m_*) because divergence events are not present. Details of this evaluation are described in the supplement. Including a hazard function that changes with time *t*, for the divergence parameters leads to a more complicated situation. The probability that a divergence event comes before a coalescent or a migration event is

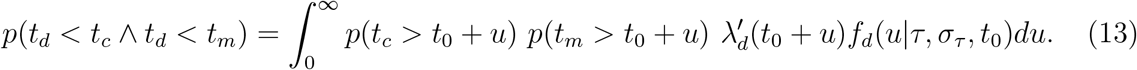

The *t_d_* = *t*_0_ + *u* is the time of a divergence event, *t_c_* is the time of a coalescent event, and *t_m_* is the time of a migration event. The 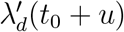 and *f_d_*(*u*|*τ*, *σ_τ_*, *t*_0_) are defined in equations 4 and 6. Since *p*(*t_c_* > *t*_0_ + *u*), *p*(*t_m_* > *t*_0_ + *u*) become *e*^−*uλc*^ and *e*^−*uλm*^, respectively, we can write

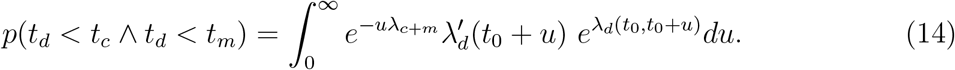

Unfortunately, the integral in Eq. (14) and its equivalents, *p*(*t_c_* < *t_d_* ∧ *t_c_* < *t_m_*) and *p*(*t_m_* < *t_c_* ∧ *t_m_* < *t_d_*), need to be solved numerically. This makes calculations very slow. The problem stems from the time-dependence of the divergence rate 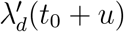. Looking for a faster way to compute these quantities, we use an approximation. Instead of solving the integral in Eq. (6) numerically, we approximate using the midpoint rule. We replaced the midpoint *t*_0_ + *u*/2 with a fixed value *t*_0_ + *ϵ* where *epsilon ≤ u*:

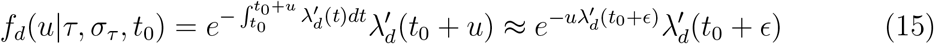

This approximation leads to a simpler formulation of Eq. (14) which now becomes:

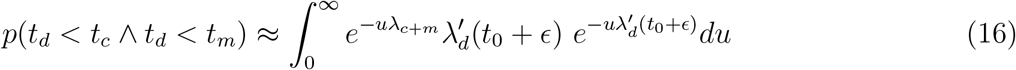

All λ*_i_* are constant with respect to *du* and therefore the integral can be solved using the substitution rule, and we get

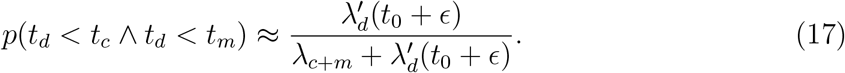

A comparison for different values of *θ*, *M*, and divergence times *τ* shows that the integral in Eq. 14 and the ratio in Eq. 17 lead to very similar values (Figure 2) This result simplifies Eq. 12 for the coalescent with divergence and migration. The general equation for all events becomes

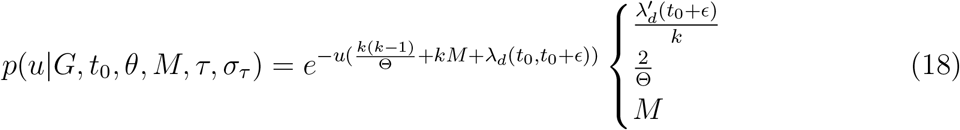

**Figure 2:**
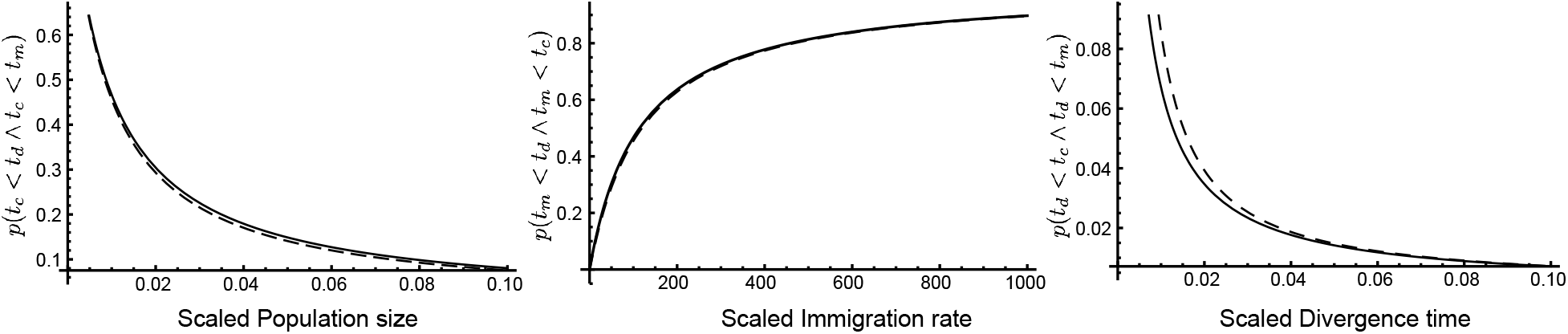
Numerical comparison of the full integral solution in Eq. 14 (dashed curves) and the midpoint approximation in Eq. 17 (black lines) of probabilities of occurrence for (from left) mutation-scaled population size *θ* (with fixed parameters *M* = 100, *τ* = 0.005, *σ_τ_* = *τ*, *t*_0_ = 0), mutation-scaled immigration rate *M* (with *θ* = 0.01, *τ* = 0.005, *σ_τ_* = *τ*, *t*_0_ = 0), and mutation-scaled divergence time *τ*(with *θ* = 0.01, *M* = 100, *σ_τ_* = *τ*, *t*_0_ = 0).

These formulas are used in Eq. 11 to calculate the probability of a genealogy given all the parameters *f*(*G*|*ρ*). The topology and the all the times of all the events are fixed in this genealogy, so we can use the time interval *u* between events to replace *ϵ*.

The exposition in the section used only one rate for each of the event-types, in reality there may be many rates for each type, for example a model with two contemporary populations, one ancestral population, and gene flow among the contemporary populations will lead to three coalescent rates, 2 immigration rates and 1 rate for the divergence time and its standard deviation.

### Implementation

The approach was implemented into the program Migrate (Beerli 2006). New parameter values were drawn from prior distributions, for example from an exponential distribution with fixed mean or a uniform distribution with a lower and upper bound. The genealogy-change proposal was described by Beerli and Felsenstein (1999), the procedure remains the same except that for the proposal of a new event and its time. In earlier versions of Migrate the time was drawn by solving Eq. 3 for the time interval *u* using a random number on the interval (0,1] as the probability, and then the probability of a particular event at that time *t*_0_ + *u* was calculated. The hazard function for the splitting rate added considerable complexity. Instead of proposing a time for any event, we chose to propose a time for each possible event independently. For example, the proposed interval u of the splitting time using Eq. 8 is

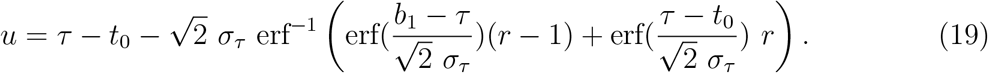

Thus, for every change of the genealogy we need to propose times for coalescence, migration, and divergence events. Among these times we pick the event with the shortest time. This approach allows us to draw the events at the correct frequency without calculating the complex ratio described in the earlier section; both, the earlier and this new calculations take about the same amount of time.

In contrast to other programs Migrate does not need a specific guide tree to specify the order of the splitting events. It uses an extension of the adjacency matrix introduced into Migrate in 2001 (Beerli and Felsenstein 2001). This matrix defines the connections among the populations by migration events and or divergence events. It can specify particular divergence models without the need to define the order of the splitting times; for example, for a model in which two island populations were colonized independently from a mainland population, Migrate does not force the user to specify an order of the time of the colonization events. We caution that our approach is not equivalent to exploring all possible population trees. Comparisons of different population trees are possible by treating each population tree as a new hypothesis and run each of these hypotheses independently, followed by Bayesian model comparison. Tutorials, source code, and executables can be found on the Migrate website (http://popgen.sc.fsu.edu).

### Simulation

Simulations were performed over a wide range of (true) divergence times *τ* from 1/512 × *N_e_* to 8.0 × *N_e_* generations. We performed three sets of simulations. The first simulation set explored the accuracy of a simple divergence model (Figure 3A). For each divergence time, four replicate datasets were simulated using the program ms (Hudson 2002) to generate the genealogies. Our own program migdata (available on www.bitbucket.com/pbeerli/popsimulate) used these genealogies to generate sequence data. To explore the effects of the number of loci, we generated datasets with 2- and 10-locus datasets for all divergence times. For a subset of divergence times, we also generated 1000-locus datasets. The second simulation set explored the interaction of immigration and divergence. We used a scenario with two populations exchanging 1 migrant every 16 generation, 4*Nm* = 0.25, and 1 migrants every 4 generations, 4*Nm* = 1.0, respectively (Fig. 3C). The immigration numbers guarantee the longterm maintenance of population structure. The third simulation set explored the effect of the estimation of the standard deviation of the divergence time. We ran simulations using our simulator speciessim.py (available on www.bitbucket.com/beerli-lab/beerlisim) using the same setting of the divergence times as before, but changed the standard deviation of the divergence time to values of *σ_τ_* = *τ*/10^4^, *σ_τ_* = *τ*/2, and *σ_τ_* = *τ* for datasets of 10 loci and compared these with the simulations of ms which simulates divergence times only with *σ_τ_* = 0.

**Figure 3:**
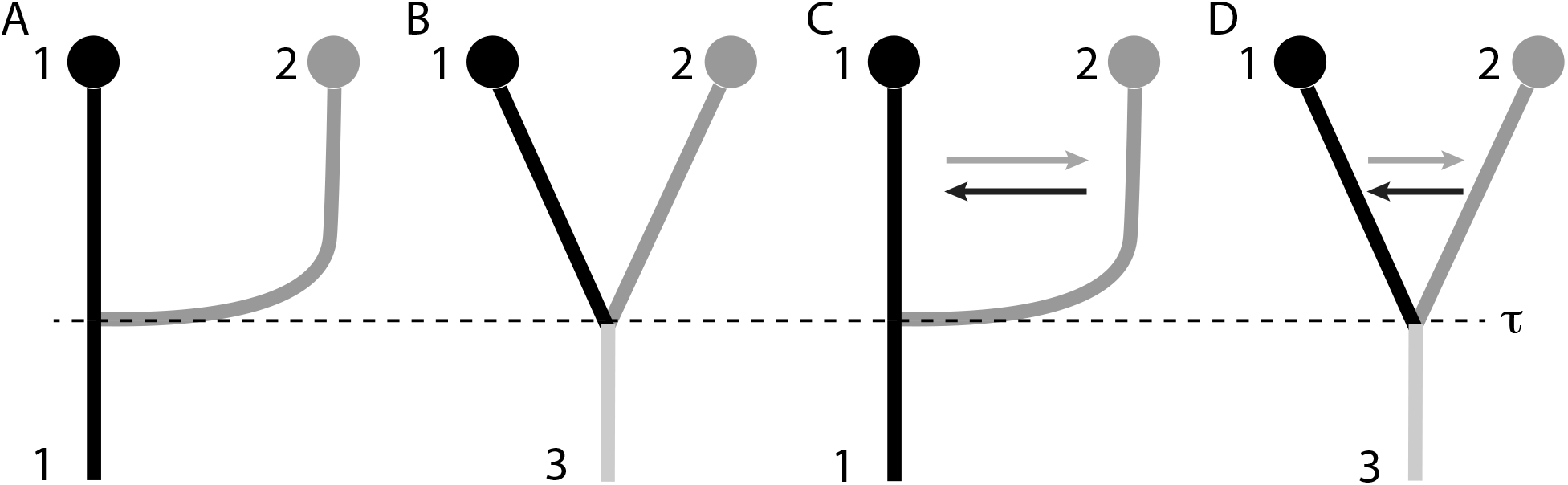
Simulation and Analysis Scenarios: (A) Population 2 splits off from population 1; (B) The ancestral population 3 splits into two contemporary populations; (C) and (D) include immigration. Models (A) and (C) were used to simulate data.

We picked an exponential prior for the mean of the divergence time distribution with an upper bound that was 3x larger then the oldest divergence time simulated; the same prior was used for all different divergence times; in pre-runs, we established that the prior choice does not change the general results. Our choice of prior and its width was made so that we could run all simulations with as little changes of options as possible. For run with real data we suggest that the range of the priors are evaluated with test runs, the divergence time *τ* in Migrate is scaled by mutations; it is on the same scale as the mutation scaled population size Θ. The choice of the prior for the standard deviation is more consequential when immigration is co-estimated, large prior ranges interact with immigration-rate estimation; consequently, we picked a small prior range for the standard deviation for the second set of simulations and also for the comparison with other programs. Simulations were run on a computer cluster with a various number of computer cores; the 10-locus simulations were run on 21 computer cores. After experimentation with run-length, we established that runs that took about 10-15 minutes are sufficient for our simulation data sets. The 1000-locus datasets were run on 40 cores and took about 50 minutes.

### Comparison with other program estimating divergence times among populations

We compared our approach with IMa2p (Sethuraman and Hey 2016), Lamarc (Kuhner 2006), and BPP (Yang and Rannala 2010) using simulated data from population models shown in Figures 3A and 3C with immigration rates of 4*Nm* = 0.0, 4*Nm* = 0.25, 4*Nm* =1.0 and divergence times of [0.0625, 0.125, 0.5, 1.0, 2.0, 4.06.0, 8.0] × *N_e_* generations. The run conditions for all programs are available in the electronic supplement and the data-converter from the Migrate format to the other programs is available from https://github.com/pbeerli/dataconverters.

### Model selection

We analyzed the simulated data used for the comparison with other programs using multiple models. Figure 3 shows the population models that were used to generate the simulated data and also models that were used to analyze the simulated data. Datasets were generated with and without immigration (Fig. 3a and 3c), we analyzed each of these datasets with all four models (Fig. 3a-d). We generated 24 datasets with no immigration and 24 datasets with 4*Nm* = 0.25 for divergence times of [0.0625, 0.125, 0.5, 1, 2, 4, 6, 8] × *N_e_* generations, this lead to 3 replicates for each divergence for each scenario. To compare and rank the models we used a Bayesian framework; we approximated the marginal likelihoods using our implementation of thermodynamic integration (Beerli and Palczewski 2010; Palczewski and Beerli 2014).

### Example using samples of complete genomes of the Zika virus

The Zika virus (ZIKV) is a pathogen distributed by mosquitoes. It was originally described in Africa. Subsequently, ZIKV was then brought to all continents via infected hosts. Gatherer and Kohl (2016) discuss the distribution routes of ZIKV based on the dates of incidences. The expansion followed this pattern: Origin in Africa, outbreaks in Asia, and then outbreaks in the Americas.

Complete ZIKV genomes from locations in Nigeria (*n* = 5), China (*n* = 12), Brazil (*n* = 13), Mexico (*n* = 2), Guatemala (*n* = 2), Panama (*n* = 4) and Puerto Rico (*n* = 2) were obtained from the NIAID Virus Pathogen Database and Analysis Resource (ViPR) (Pickett et al. 2012) through the web site at http://www.viprbrc.org/. The sequences were aligned with MUSCLE (Edgar 2004, aligned dataset in electronic supplement). We pooled the locations Mexico, Guatemala, Panama, and Puerto Rico for the analysis.

We then explored four different main population models (Fig. 4). The model group *a* specified the expansion from Africa to Asia to Brazil to Central America. The model group *b* specifies the expansion from Africa to Asia and, independently, from Africa to Brazil to Central America. The model group *c* is a hybrid of models *a* and *b* where one population is the admixture of two populations. We considered the Brazilian lineages a potentially admixed population from African and Asian lineages. Occurrences of ZIKV in Central Americas and Brazil were reported contemporaneously, suggesting that there may be not enough mutations to separate Brazilian and Central American lineages; we combined the samples from Brazil and Central America (model group *d*); otherwise, *d* is equivalent to *c*. The variants *a*′, *b*′, *c*′, and *d*′ include recurrent immigration from the source populations.

**Figure 4:**
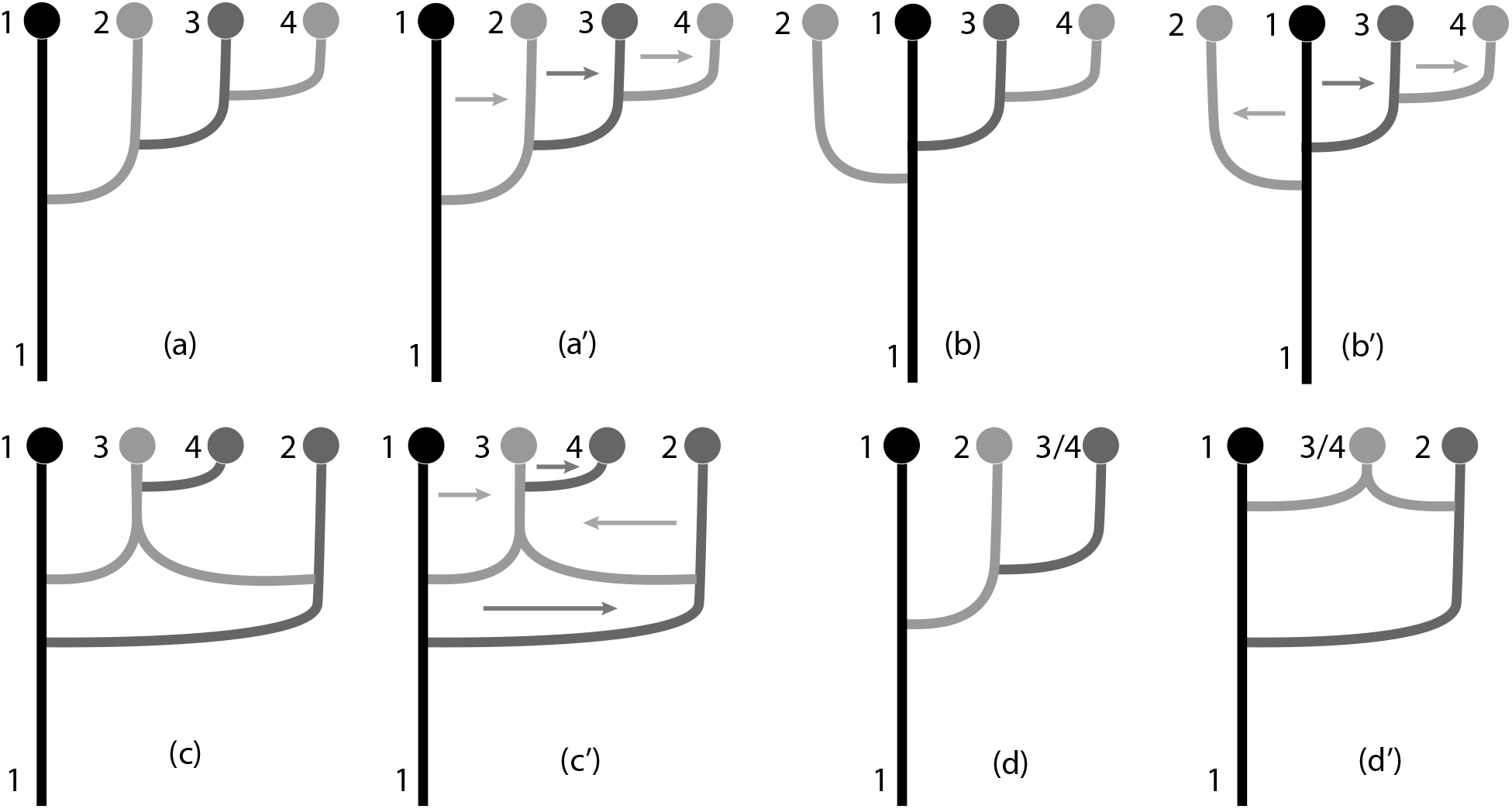
Eight population models used to analyze the ZIKV dataset. Tip labels are 1=Africa, 2=Asia, 3=Brazil, 4=Central America, and 3/4 = Brazil and Central America combined. The arrows mark migration directions.

## Results

### Simulation

#### Splitting time estimation

The inference code was evaluated using simulations over a wide range of divergence times and two different migration rates after the population split.

Figure 5 compares the estimated population split time 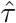 with the population split time *τ* used to simulate the data without migration. The estimates track the simulated split times well, although the estimates of large divergence times are underestimated. As expected, the estimates from two-locus data sets show more spread than those from ten-locus or 1000-locus datasets. A comparison of the 95% credibility intervals of runs with 2, 10, and 1000 loci shows this trend: standardizing the credibility interval with the observed mode 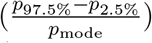 leads to averages of 6.88 for two loci (*N* = 271, *σ_τ_* = 14.01), 2.45 for ten loci (*N* = 280, *σ_τ_* = 5.58), and 0.37 for 1000 loci (*N* = 28, *σ_τ_* = 0.72).

**Figure 5:**
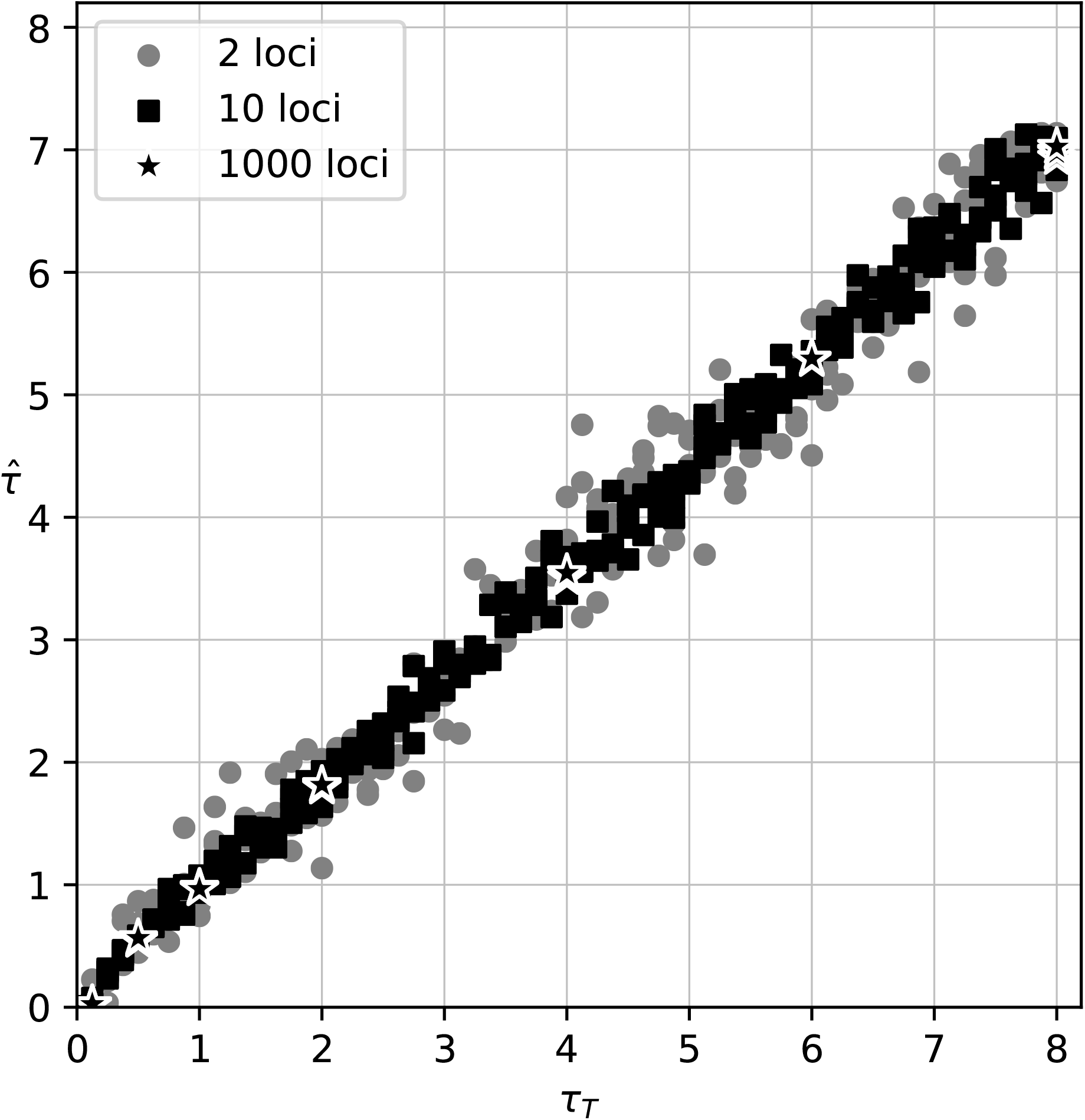
Comparison of estimated divergence time 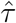 with the true divergence time *τ_T_* that was used to simulate 2-locus, 10-locus, and 1000-locus data. The data were simulated and analyzed using the model shown in Fig. 3A. Units of 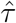 and *τ_T_* are in *N_e_*× generations.

#### Splitting time estimation under the isolation with migration model

The simulation results with migration deliver a more complicated message. Simulations with low recurrent immigration rates (4*Nm* = 0.25) during the time interval from today to the population split track the true population split often quite well but has a considerable fraction of runs that underestimate the divergence time (Fig. 6). With a higher immigration rate (4*Nm* = 1.0) 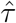 underestimates the true divergence time of datasets that were created using a high divergence time and overestimates the divergence time for very recent divergences.

**Figure 6:**
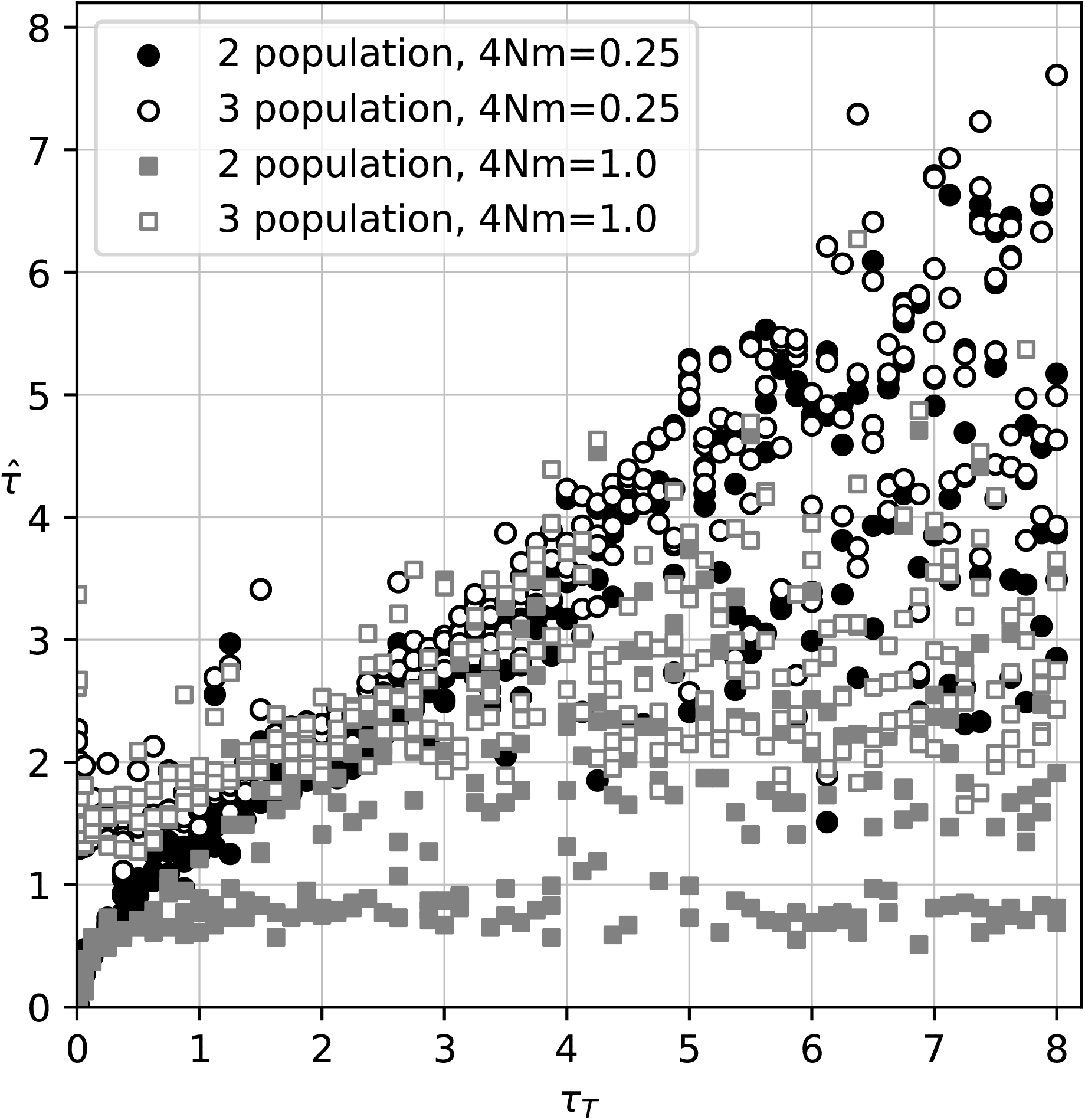
Comparison of estimated divergence time 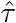 and the true *τ_T_*. The data were simulated using the model shown in Figure 3C, and analyzed using models Figure 3C and 3D. The number of immigrants per generation were 4*Nm* = 0.25 and 4*Nm* =1.0, respectively. Units of 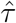 and *τ_T_* are in *N_e_*× generations.

**Figure 7:**
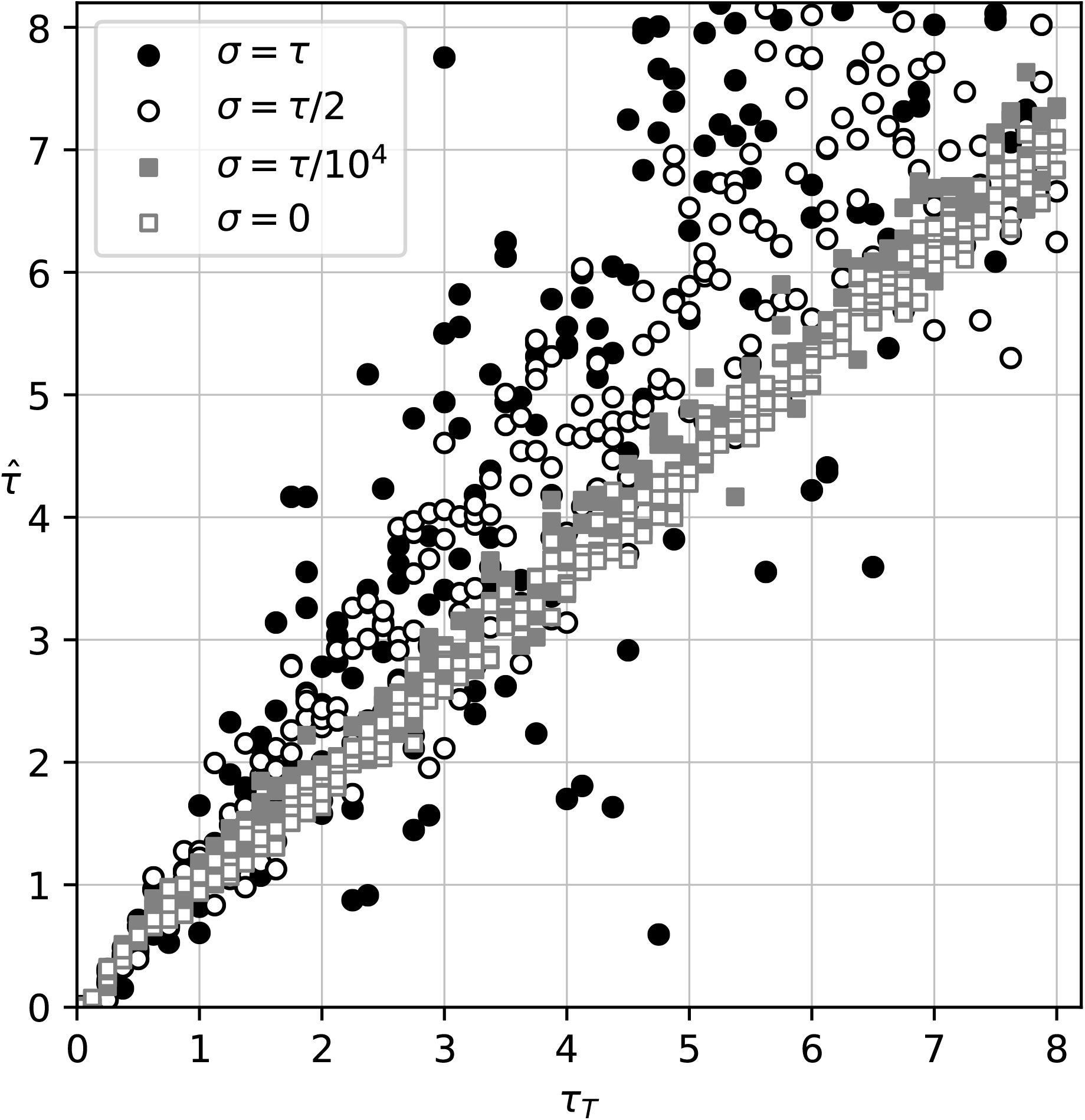
Comparison of estimated divergence time 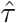 and the true *τ*. The 10-locus data were simulated without immigration but with four different standard deviations for the splitting time (*σ_τ_* = *τ*, *σ_τ_* = *τ*/2, *σ_τ_* = *τ*/10^4^, and *σ_τ_* = 0).

#### Effect of simulated uncertainty of splitting time

Our approach allows the estimation of the splitting time and the standard deviation of the splitting time. All current simulation methods, except our own speciessim.py, assume a defined time when the ancestral population splits into offspring populations. Our simulation and estimation model allow uncertainty about this time, a small standard deviation *σ_τ_* = 1/10000 will result in simulated data sets that mimic the standard simulation method in MS. Large standard deviation leads to datasets with skewed distributions of divergence times because the divergence time cannot be negative (looking backward in time), and such times had to be redrawn to generate the simulated genealogies. About 15.8% of all random draws from a Normal distribution will be smaller than *τ* − *σ_τ_*. This resulted in datasets that come from older divergence times on average and will result in higher divergence time estimates than the divergence time *τ* used to generate the datasets.

#### Exploration of the splitting time bias in the IM model

If the immigration rate is high, population divergences that happened far in the past are problematic to estimate because in comparisons (Fig. 6) we detect a bias towards more recent split time estimates than those simulated. To investigate this bias, we have simulated genealogies with sample sizes of 40 and 100 with the same parameters used to create the data reported in Figure 6 and recorded the number of lineages present at the time of population split (70 time points between split times of *τ* from 1/512 × *N_e_* to 8.0 × *N_e_* generations. Figure 8 reports the percentage of datasets that have two or more lineages available at these 70 time-points (*N*=1000 for each time point). The graphs for 4*Nm* = 0.25 (low) and 4*Nm* = 1.0 (high) differ starkly in the percentages with high divergence times. With high immigration rates (4*Nm* =1.0) the chance of having the sample coalesced to a single lineage increases considerably, for example, fewer than 20% of the datasets have information about a divergence time of 4*N_e_* generations. Once a sample coalesces into a single lineage all information about the historical processes is lost, and any inferred result will only come from the prior and is independent of the data. Increasing the number of individuals from 40 to 100 for each dataset does not improve the number of available lineages at the divergence time. With low immigration rates, the time of the most recent common ancestor is beyond the divergence time. Thus, the remaining lineages may have information about the splitting time.

**Figure 8:**
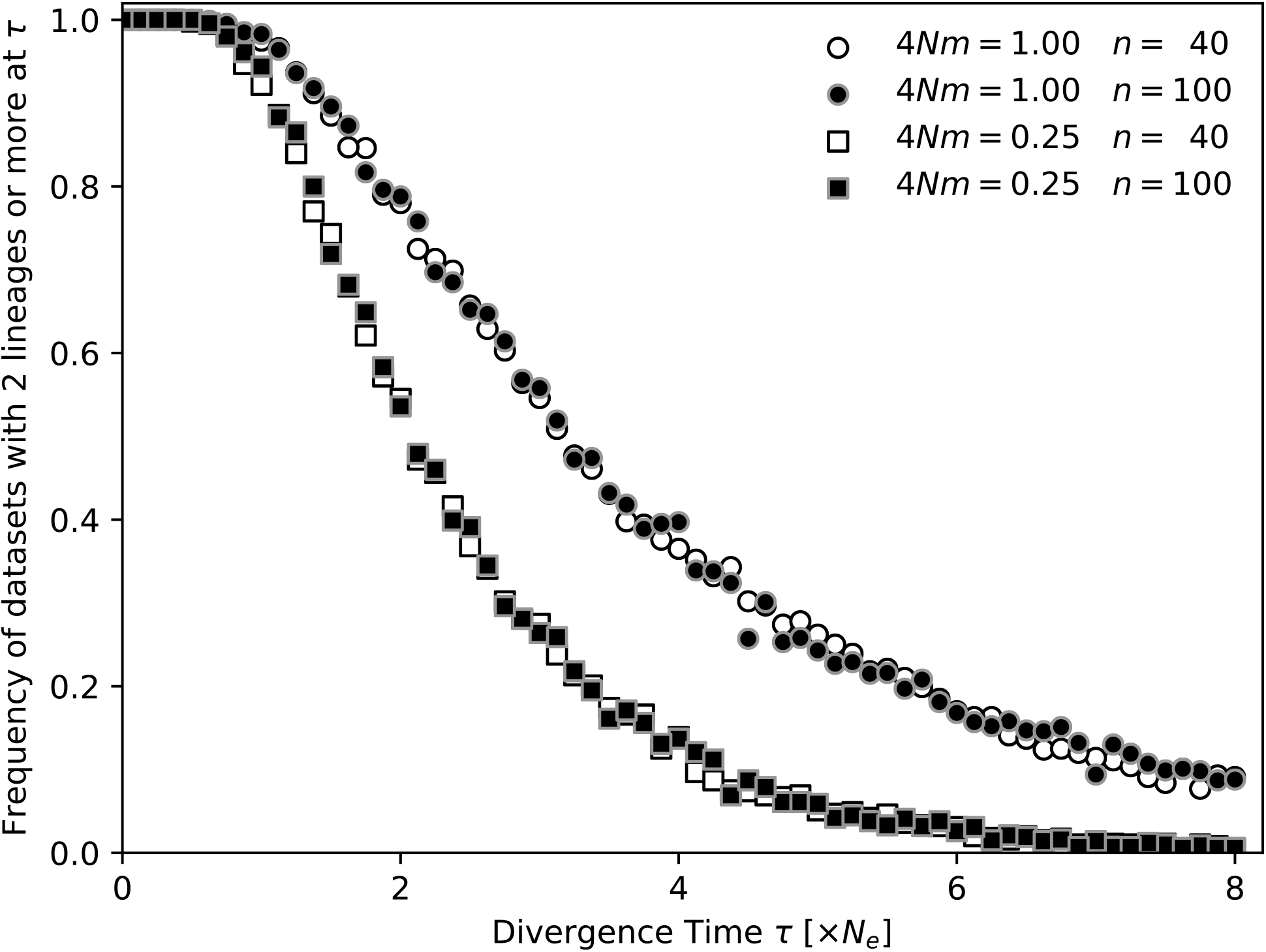
Percentages of simulated datasets with two or more lineages in the sample at divergence time *τ*. For each divergence time *τ* 1000 datasets were simulated.

#### Comparison with other programs

Three sets of simulated data for immigration rates of 4*Nm* = [0.0, 0.25,1.0] were used to compare the results of four different programs: Migrate, IMa2p, Lamarc, and BPP. Figure 9 shows the results for these comparisons. We tried to spend similar efforts for all programs: IMa2p was run on 4 cores for 4 hours, Lamarc was run for approximately 3 hours, BPP was run for about 1.5 hours, Migrate was run for 10-15 minutes on 20 compute cores; Lamarc reported the most variable results, looking at their outputs the runs may have struggled to converge. Divergence times can be well estimated by all programs when recurrent gene flow is zero and the true divergence time is smaller than 2*N_e_* generations. All programs show a bias when the true divergence times become large compared to the population size of the sampled populations, Migrate shows a smaller bias than the others. Results become more unpredictable when gene flow is larger than zero. BPP underestimates the divergence time with any immigration, but it is the only program that does not estimate immigration rates and therefore all immigration events will contribute to the estimation of divergence time. Migrate and Lamarc overestimate the divergence time when the true divergence time is low, but Migrate estimates larger divergence times with immigration well. IMa2p estimates low divergence times with low immigration rates well, but divergence time estimates are underestimated with large true divergence times. With immigration rates of 4*Nm* = 1.0 all programs fail to estimate accurate divergence times. Migrate and IMa2p deliver very similar results. Both are overestimating divergence times when the true divergence is small and underestimating when the true divergence times are large.

**Figure 9:**
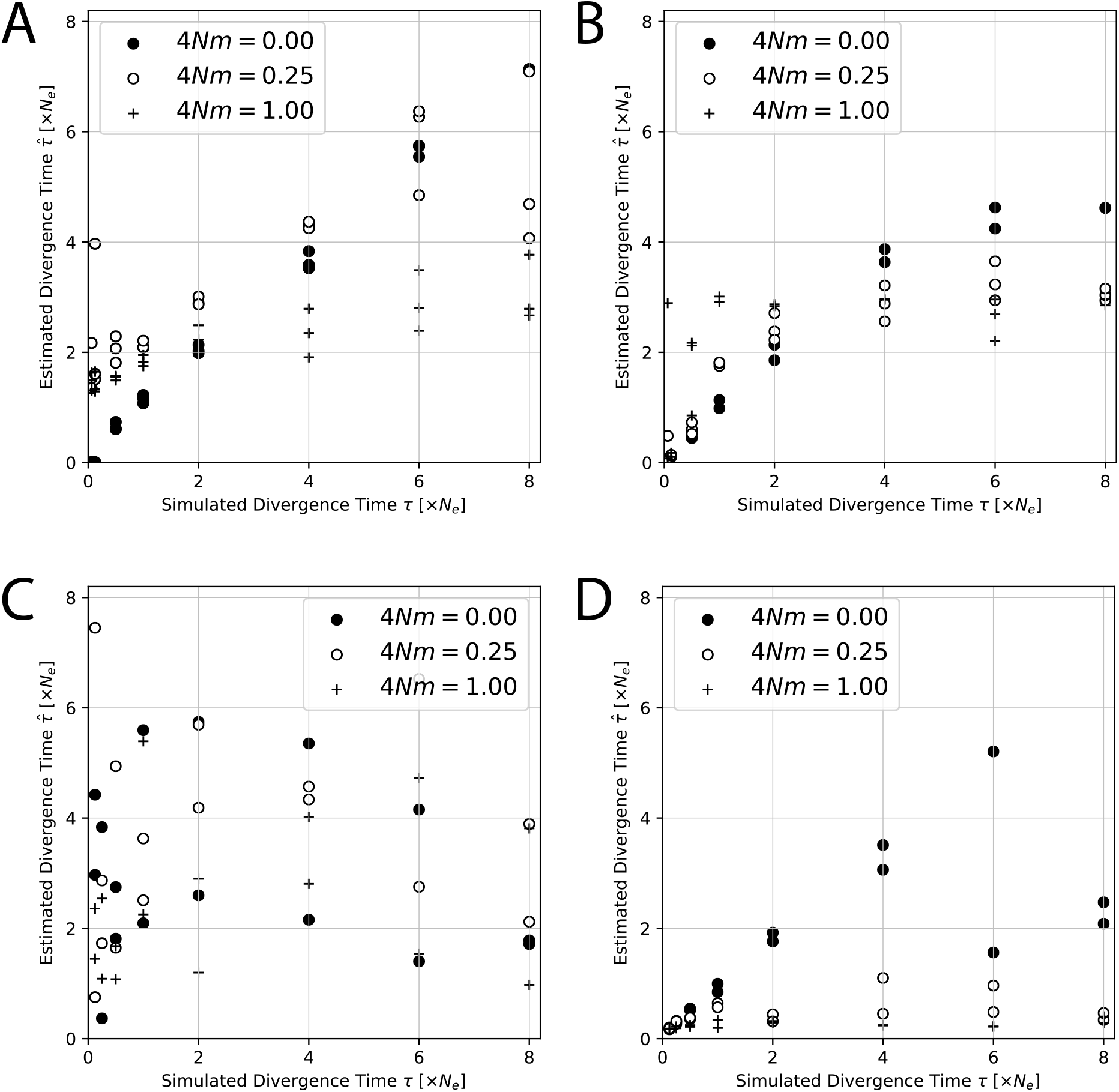
Comparison of estimated divergence time 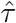 and the true *τ_T_* for (A) Migrate, (B) IMa2p, (C) Lamarc, (D) BPP. The data were simulated using the model shown in Figure 3C, and analyzed using models Figure 3C and 3D. The number of immigrants per generation were 4*Nm* = 0.0, 4*Nm* = 0.25, and 4*Nm* = 1.0, respectively.

#### Model selection

We compared four models: a two-population and a three-population isolation-with-migration (IM, Fig. 3C, D) and a two-population and a three-population isolation-only model (IO, Fig. 3A, B) for these divergence times: [0.0625, 0.125, 0.5, 1.0, 2.0, 4.0, 6.0, 8.0] × *N_e_*. Datasets simulated with the two-population IO model preferred the two-population IO model for divergence times larger or equal to 0.125 × *N_e_* generations (19 out of 24 tests); two of the three replicates with divergence time of 0.125 × *N_e_* generations and the replicates with divergence of 0.0625 × *N_e_* generations preferred the IM model (5 out of 24 tests). When the data was simulated with the two-population IM model with 4*Nm* = 0.25 (Fig. 3C) and tested against the four models, 24 out of 24 comparisons preferred the 3-population IM model. All model comparisons are reported in the electronic supplement.

### Zika virus dispersal

Table 1 shows the model probabilities and the log marginal likelihoods for different models (see Fig. 4) for the Zika virus (ZIKV). The model in which the expansion followed a route to the east from Africa is the most likely model. The best model is a simple colonization model without migration. Figure 10 shows the population tree of the best model with mutation-scaled population sizes and divergence times. Yokoyama and Starmer (2017) used an estimate of the mutation rate for various lineages of the Asian ZIKV lineages, suggesting that the mutation rate per year has accelerated and is *τ* = 0.004/year, but can be as low as 0.0005/year. Faria et al. (2016) estimate a phylogenetically derived mutation rate of *τ* = 0.00098 to 0.00106 per year. Migrate estimated the mutation-scaled divergence time 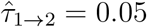 and 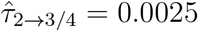 assuming that the mutation rate is per generation. We did not find any clear characterization of generation time for ZIKV in the literature. We equate generation time here as successful transmissions among hosts per year and not the number of replications of an individual ZIKV within a host. Early records from Africa date to 1947 and early records from Asia date to 1951. Thus, gene flow of ZIKV from Africa to Asia was most likely around 1950. The ZIKV outbreak in Brazil started in 2015 (Faria et al. 2016). Ignoring the precise sampling dates and assuming the divergences were 67 and 3 years ago then we calculate about 5 generations per year (67/(0.05/0.004) = 5.2 and 3/(0.0025/0.004 = 4.8) using the high mutation rate. The lower mutation rate (~ 0.001) would lead to 1.34 and 1.2 generations per year, respectively.

**Table 1:** Log marginal likelihoods and model probabilities of biogeographic models: (a) eastward, (b) westward, (c) admixture, and (d) three-population models (Fig. 4) and Zika viruses. The → mark colonizations, the ⇒ mark colonizations with recurrent immigration. Numbers are population labels: 1=Africa, 2= Asia, 3=Brazil, 4=Central America.

| Model |  | Zika |  |  |  |
| --- | --- | --- | --- | --- | --- |
| | | $\ln(\text{mL})$ | LBF* | Prob. | Rank |
| a | $1 \rightarrow 2 \rightarrow 3 \rightarrow 4$ | -25762.13 | -95.61 | 0.0 | 3 |
| a' | $1 \Rightarrow 2 \Rightarrow 3 \Rightarrow 4$ | -26078.46 | -411.94 | 0.0 | 6 |
| b | $2 \leftarrow 1 \rightarrow 3 \rightarrow 4$ | -25824.41 | -157.89 | 0.0 | 5 |
| b' | $2 \Leftarrow 1 \Rightarrow 3 \Rightarrow 4$ | -26121.92 | -455.40 | 0.0 | 8 |
| c | $1 \rightarrow 3 \leftarrow 2 \leftarrow 1, 3 \rightarrow 4$ | -25786.83 | -120.31 | 0.0 | 4 |
| c' | $1 \Rightarrow 3 \Leftarrow 2 \Leftarrow 1, 3 \Rightarrow 4$ | -26099.01 | -432.49 | 0.0 | 7 |
| d | $1 \rightarrow 2 \rightarrow 3/4$ | -25666.52 | 0.0 | 1.0 | 1 |
| d' | $1 \rightarrow 3/4 \leftarrow 2$ | -25703.50 | -36.98 | 0.0 | 2 |

\* LBF = $\ln$ Bayes factor against the best model $d$

**Figure 10:**
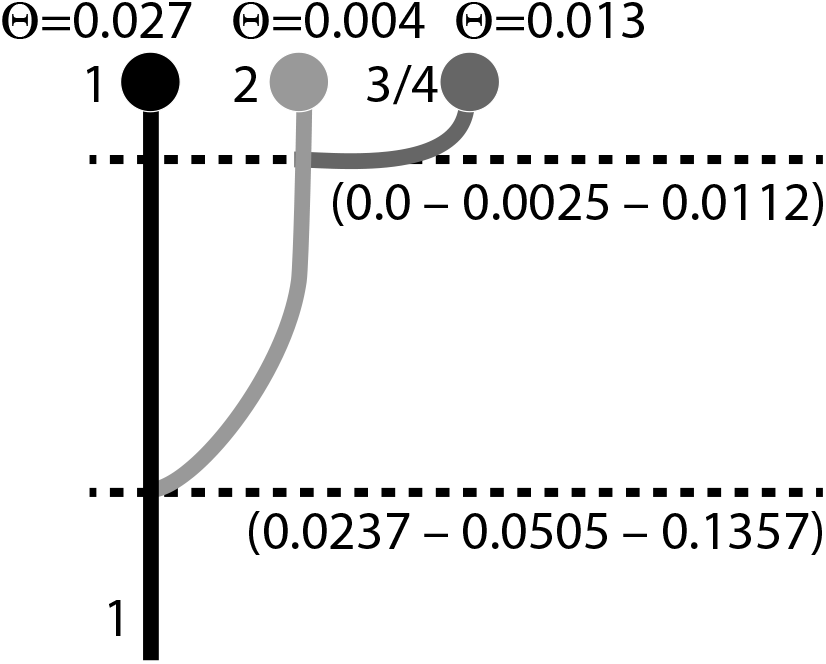
Mode and 50% credibility intervals of the splitting times and population sizes of the best model for Zika virus (model *d*; Fig. 4)

## Discussion

We have developed a model to incorporate population splitting and population admixture. Our algorithm differs from other algorithms because we treat the splitting times as random variables with truncated Normal distributions. This method allows a wide range of analyses, such as having populations split from an ancestral population or having population split from a population that is the same today and in the past.

The joint estimation of divergence time and population sizes without immigration from large genetic datasets seems feasible with little error. Our simulations assumed informative loci and no complications with the finite mutation model. However, Migrate can handle site rate variation and more complex mutation models than the F84 model used in the simulations. We assume that Migrate has similar vulnerabilities as IM when tested with deviations of the model (cf. Strasburg and Rieseberg 2010).

Migrate runs each locus as an independent unit and thus can efficiently run large datasets, such as the 1000 loci datasets used in the simulations, in reasonable time on a cluster computer: the 1000-locus datasets for Figure 5 were run on 40 cores and finished under an hour. The comparisons of the data with 2, 10, 1000 loci show that with informative loci, we may not need to have many loci to extract the most likely parameter value, although the variances of the 1000 locus runs are smaller than the others.

It seems straightforward to use an immigration with divergence model (IM; Nielsen and Slatkin 2000), but little exploration about the power of the inference has been conducted. Strasburg and Rieseberg (2010) highlighted that assumption misspecification can lead the program IM (Hey 2010) to deliver biased answers. Recently, Quinzin et al.(2015) evaluated the program IM and observed that divergence time estimates are more accurate if migration is low and if the populations are large compared to the divergence time. We find similar patterns with Migrate and IMa2p. Additionally, our simulations of the immigration with migration model show a deeper problem with such inferences, even when assumptions are met. Looking backward in time, once all samples have coalesced, no information is left to estimate parameters. In a model with immigration and population splitting there has to be a balance so that we can see the effects of one or the other: if the migration rates are small, then all sample lineages, looking backward in time, will have joined the ancestral population before having experienced a migration event. In contrast, with high immigration rates, it becomes very likely that all lineages have coalesced into one lineage before the expected splitting time. Figure 8 shows that many sample data never experience a population split. It will undoubtedly be difficult to estimate an event that did not leave a trace in the sample. Hence, the estimated divergence times will not reflect the true splitting time and will be too close to the sampling date. However, with small immigration rates, it is possible to recover splitting times that are further in the past (Fig. 6). The same simulations also show that it is unproblematic to estimate splitting times that are old when there is no immigration. Our results for Migrate, IMa2p, Lamarc, and BPP suggest that one should use caution when using models with immigration and population splitting times. This dependency is independent of the estimation method, and certainly will include other than the tested methods, too. We believe that this dependency has and will lead to incorrect reports of divergence times: divergence times are reported to be more recent than they are.

The direction of the expansion of the pathogens as estimated by Migrate is simply based on genetic data and coalescence-based population genetic models. The used data are not very informative because only a few sequences from Africa are present. This does not allow to pinpoint the expansion from Africa to Asia with good precision. It is also likely that the expansion from Brazil to Mexico and other countries is not very informative because only a few sequences from a large area (Mexico, Guatemala, Panama, and Puerto Rico) were used. Gatherer and Kohl (2016) summarized the literature on the spread of the Zika virus and shows an expansion from Africa to Asia to Pacific islands and then to South America. They used incidences of confirmed Zika virus infections and already published phylogenetic trees to report a map of its spread. It is comforting that our population genetics approach recovers the same paths as the more detailed historical records of infections; genetic data will be particularly useful for pathogens for which we may not have detailed incidence records.

We have presented an alternative to current estimations of divergence time among populations. Our method not only allows considering the splitting times but even allows to date admixture of a population from two or more ancestral populations. The simulations suggest that fairly variable data are needed. Estimation of splitting times alone is robust over a wide range of simulated splitting times, whereas models that allow migration and splitting times (IM model) simultaneously suffer considerable difficulties estimating splitting times that are far in the past when population sizes are small, and immigration rates are high. These difficulties are caused by the sparsity of lineages far in the past, a situation that is well known (Heled and Drummond 2008). Improving these estimates will depend on the number of loci, the number of individuals, and data with different sampling dates.

## Acknowledgement

We thank Thomas Uzzell for giving us suggestions to an earlier manuscript. Suggestions by the associate editor and an anonymous reviewer helped greatly to improve this work. The staff at the Research Computing Center at Florida State University helped us to run our simulations smoothly on the high performance computing cluster.

## Author contributions

MP and PB developed the idea and algorithms, SM improved the algorithms. MP wrote a test version of the method. PB incorporated the method into the program Migrate. HA wrote a simulator program to test the program. PB run all analyses in this study. PB, MP, and SM wrote the manuscript.

## Supplementary material

Simulated data, aligned real data, and the elaboration on some of the equations are available from http://github.com/pbeerli. The software Migrate is available at the Migrate website http://popgen.sc.fsu.edu, simulation software is available at http://github.com/pbeerli.

## Supplement

### Constant λ_1_ and λ_2_

In the standard structured coalescence with two forces, genetic drift and recurrent gene flow between isolated populations we have two different types of rates, one type for coalescences and another type for immigration events. Both types can be considered exponential rates; there is a constant risk over time that one or the other event happens, so we can calculate

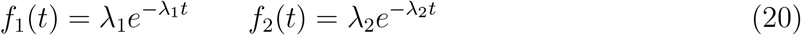

We consider first the case where the event with *λ*_1_ happens first (*T*_1_ < *T*_2_). We find *P*(*T*_1_ < *T*_2_) assuming that

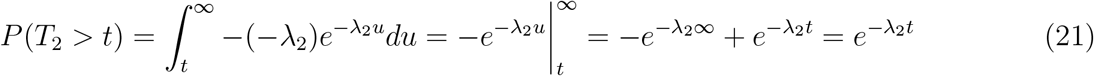

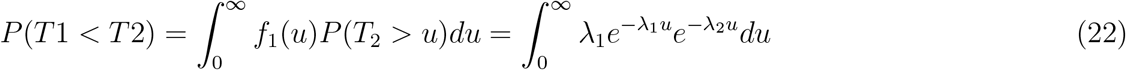

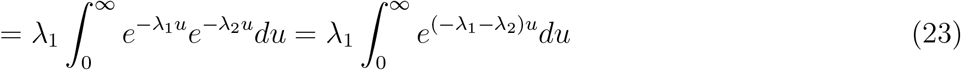

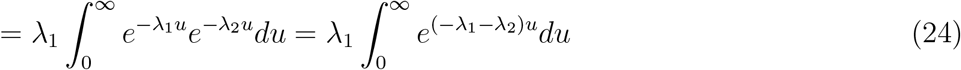

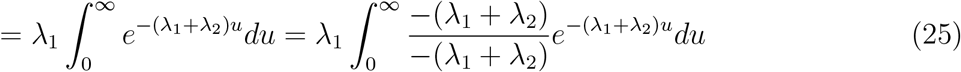

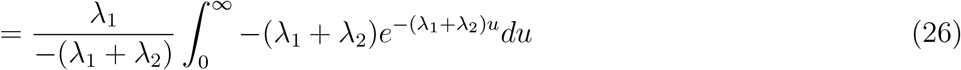

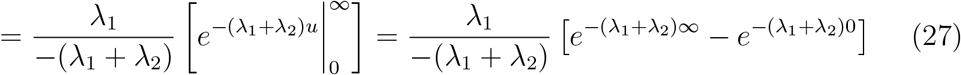

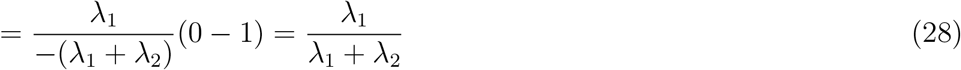

### Non-constant and constant rates

In this section, we consider three different cases. Suppose we have three events

1. The divergence event 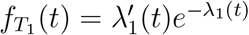, Time related to it call *T*_1_
2. The coalescent event → *f*_*T*_2__(*t*) = *λ*_2_*te*^−*λ*_2_*t*^, Time related to it call *T*_2_
3. The Migration event → *f*_*T*_3__(*t*) = *λ*_3_*te*^−*λ*_3_*t*^, Time related to it call *T*_3_

Divergence has a rate that changes with the time, the risk of switching increases the longer we wait and is non-constant, This leads to complication in finding a solution to the integral.

### Divergence happen first

We need to find *P*(*T*_1_ < *T*_2_&*T*_1_ < *T*_3_). We know

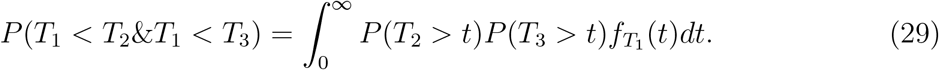

Fist we find *P*(*T*_2_ > *t*) and *P*(*T*_3_ > *t*) as

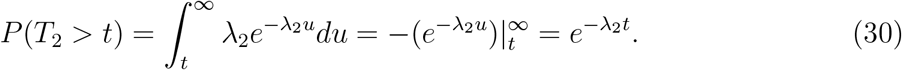

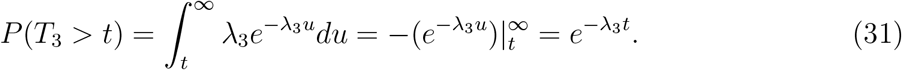

Using Eqs. (29)-(31) we have

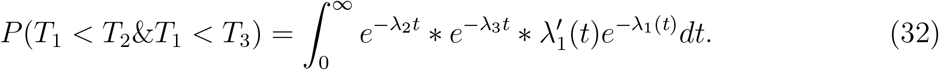

### Coalescent happen first

We need to find *P*(*T*_2_ < *T*_1_&*T*_2_ < *T*_3_). We know

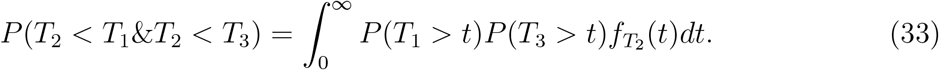

Fist we find *P*(*T*_1_ > *t*) and *P*(*T*_3_ > *t*) as

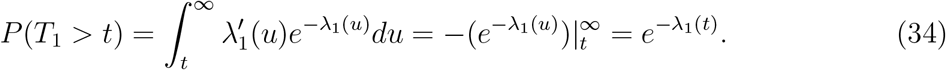

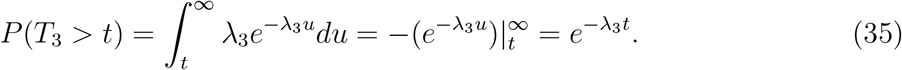

Using Eqs. (33)-(35) we have

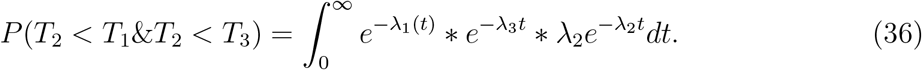

### Migration happen first

We need to find *P*(*T*_3_ < *T*_1_&*T*_3_ < *T*_2_). We know

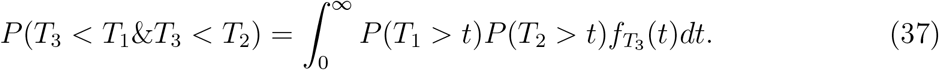

Fist we find *P*(*T*_1_ > *t*) and *P*(*T*_2_ > *t*) as

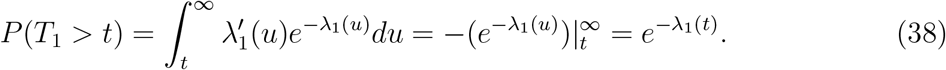

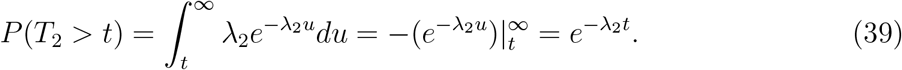

Using Eqs. (37)-(39) we have

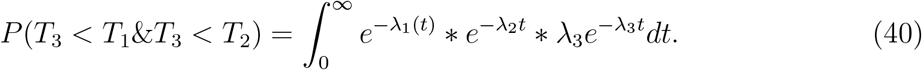

### Distribution of the time to the first event

If we have three events, Divergence *T*_1_, Coalescent *T*_2_ and Migration *T*_3_, the distribution of time to the first event *T* = *min*(*T*_1_, *T*_2_, *T*_3_) is as

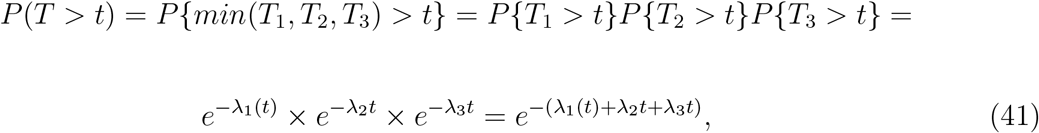

so *T* = *min*(*T*_1_, *T*_2_, *T*_3_) has an exponential distribution.

